# Trunk neural crest migratory position and asymmetric division predict terminal differentiation

**DOI:** 10.1101/2022.02.23.481590

**Authors:** Zain Alhashem, Karen Camargo-Sosa, Robert N Kelsh, Claudia Linker

## Abstract

The generation of complex structures during embryogenesis requires the controlled migration and differentiation of cells from distant origins. How migration and differentiation are coordinated and impact each other to form functional structures is not fully understood. Neural crest cells migrate extensively giving rise to many cell types. In the trunk, neural crest migrate collectively forming chains comprised of cells with distinct migratory identities: one leader cell at the front of the group directs migration, while followers track the leader forming the body of the chain. Herein we analysed the relationship between trunk neural crest migratory identity and terminal differentiation. We found that trunk neural crest migration and fate allocation is coherent. Leader cells that initiate movement give rise to the most distal derivativities. Interestingly, the asymmetric division of leaders separates migratory identity and fate. The distal daughter cell retains the leader identity and clonally forms the Sympathetic Ganglia. The proximal sibling migrates as a follower and gives rise to Schwann cells. The sympathetic neuron transcription factor *phox2bb* is strongly expressed by leaders from early stages of migration, suggesting that specification and migration occur concomitantly and in coordination. Followers divide symmetrically and their fate correlates with their position in the chain.

## Introduction

The tight coordination of cell migration and differentiation is fundamental for organ formation during development, as physiologically functional units are often composed of cells that originate at distant locations. The neural crest (NC) is a transient population of cells that arise early in development, migrate extensively and differentiate into a plethora of derivatives (Le Douarin and Kalcheim, 1999). Hence, the NC population constitutes an excellent model system to understand the interplay between migration and differentiation. Trunk NC cells (TNC) give rise to pigment cells, neurons, glia including Schwann cells, and chromaffin cells, with each migrating along stereotypic pathways (Raible and Eisen, 1994). Here we focus on the TNC migrating medially between the somitic tissue and the neural tube, the medial path in zebrafish, equivalent to the ventral pathway in chick (Raible et al., 1992), which differentiate into glia and neurons of the sympathetic and dorsal root ganglia, and melanocytes and iridophores. TNC migrating on this medial pathway form single file chains that require the establishment of two distinct migratory identities (Richardson et al., 2016): the leader cells that initiate movement and remain at the front of the group throughout migration, are the only cells capable of instructing directionality to the group. In contrast, follower cells track the leader’s movement forming the body of the chain, and require cell-cell contact for migration. Interestingly, leader and follower cells not only differ in migratory behaviour but also in morphology and cell cycle progression. Leader cells are larger, more elongated and polarized than followers. Leaders and followers present a similar total cell cycle span, but leaders present a long S-phase while followers spend most of the cycle in G_1_ (Alhashem et al., 2021a). Together these features distinguish leaders from followers, and allow the collective migration of the group, but whether these traits impact terminal TNC differentiation is unknown.

TNC migrating into the ventral pathway that halt movement lateral to the dorsal aorta, the ventral-most target, form the Sympathetic Chain Ganglia (SCG). Cells arresting migration at more dorsal positions envelope motor axons and differentiate as Schwann cells (SCs), while TNC that migrate up to, or just ventral to, the neural tube form the neurons and glia of the dorsal root ganglia (DRG; Raible and Eisen, 1994; Raible et al., 1992). How do TNC know what cell type to form? Are these a heterogenous group of predetermined cells or a homogenous naïve population? And does the collective migration of this group have any consequences for the differentiation outcome of cells? A predominant view is that NC are multipotent, with each cell able to generate multiple cell-types according to local-cues (Baggiolini et al., 2015; Douarin and Teillet, 1974; Fraser and Bronner-Fraser, 1991; Henion and Weston, 1997; Le Lievre et al., 1980; Rothman et al., 1990; Sieber-Blum and Cohen, 1980; Weston, 1991). In accordance with this model studies in mouse have shown that some NC retain stem cell capacities (Stemple and Anderson, 1993). A key question is whether individual NC undergo progressive fate restriction, whereby they lose the capacity to generate some or many fates (Kelsh et al., 2021; Le Douarin, 1986; Weston, 1991). Studies in zebrafish and avian embryos provide evidence that early migrating NC might already be a heterogeneous pre-determined population in which each cell is only capable of generating a single or a small number of cell-types, with *in vitro* and *in vivo* studies showing that the majority of NC consist of lineage-restricted cells soon after their emergence from the neural tube (Henion and Weston, 1997; Raible and Eisen, 1994). Moreover, detailed fate map in chick embryos support these results, showing that the relative position of TNC in the premigratory area, as well as the time of migration initiation is predictive of their target location and differentiation outcome (Krispin et al., 2010). However, whether TNC migratory dynamics plays a role in their differentiation outcome of has not been addressed. To analyse this question, we tracked single zebrafish TNC *in vivo* from the initiation of migration to their final target sites and defined the relationship between migratory identity and fate. Our results show that the majority of the cells in the migratory chain give rise to single type of derivatives. Moreover, the migratory identity of the cells strongly correlates with their terminal differentiation. Leader cells divide asymmetrically, giving rise to a distal daughter that retains the leader’s identity and clonally forms the SCG. The proximal sibling on the other hand, becomes a follower and differentiates into SCs. Interestingly, leader cells present an enriched expression of the sympathetic transcription factor *phox2bb* from early stages of development, suggesting that fate specification and migration occur concomitantly and in coordination. Unlike other followers, which only incorporate into the DRG, the first follower cell can incorporate into the DRG or differentiate as SCs.

## Materials and Methods

### Resource and Materials availability

Further information and requests for resources and reagents should be directed to and will be fulfilled by the lead contact, Claudia Linker

### Zebrafish husbandry and transgenic lines

Zebrafish were maintained in accordance with the UK Home Office regulations UK Animals (Scientific Procedures) Act 1986, amended in 2013 under project license P70880F4C. Embryos were obtained from the following strains: Sox10:mG, Tg(−4.9sox10: Hsa.HIST1H2BJ-mCherry-2A-GLYPI-EGFP); Tg(h2afva:h2a-GFP)kca13; Neurog1:GFP; Tg(Sox10:GFP)ba4. Offspring of the required genotypes were staged according to Kimmel et al. (1995) and selected based on health and the expression of fluorescent reporters when appropriate. Embryos were split randomly between experimental groups and maintained at 28.5°C in E3 medium.

### Live Imaging and tracking

Imaging and analysis were carried as in (Alhashem et al., 2021b). In short, embryos were mounted in 0.6% low melting point agarose/E3 medium plus 40 μM Tricaine. For long term movies, tiles of segments 6-22 were imaged in lateral views every 20 or 30 minutes from 16hpf for 3 days in an upright PerkinElmer Ultraview Vox system using 40X water immersion objective. In general, 70 μm z-stacks with 2 μm z-steps were obtained, however, 1 μm z-steps were obtained for cell volume rendering and calculation. For leaders and followers division, movies were acquired every 5 minutes. Image stacks were corrected for drift using the Correct 3D Drift Fiji plugin and 3D single cell tracking were performed with the View5D Fiji plugin. Tracks overlays were drawn using the MTrackJ and Manual Tracking Fiji plugins. Cell volume rendering and calculations were done in Imaris (Bitplane). Cell area measurements were done in Fiji using the freehand selection tool. Division angles were measured using the angle tool in Fiji. Cell speed measurements were calculated from 3D tracks according to ((SQRT((X1-X2)^2+(Y1-Y2)^2+(Z1-Z2)^2))/T)*60. Where X, Y and Z are the physical coordinates and T is the time-step between time-lapse frames. Cell directionality measurements were calculated using previously published Excel macros (Gorelik and Gautreau, 2014).

### In situ hybridisation and immunostaining

Anesthetised embryos were fixed in 4%PFA overnight at 4 °C. PFA was removed and 100% methanol was directly added. Samples were stored at −20 °C until processed. We used the RNAscope Multiplex Fluorescent kit V2 (Bio-techne, Cat No. 323110) following the manufacturer’s protocol with some modifications. Methanol was removed from samples and air-dried for 30 min at RT. 50 ml of Proteinase Plus was added for 10 min at RT and washed with 0.01% PBS-Tween for 5 min x3. Samples were incubated in 50 μl of hydrogen peroxide for 7 minutes at RT followed by 3 washes with RNAse free water for 5 min each. Samples were incubated overnight with diluted probes (1:100). Probes were recovered and samples were washed in 0.2X SCCT for 10 min x2. We followed the manufacturer instructions for AMP 1-3 and HRP C1-C4 using 2 drops of each solution, 100 μl of Opal 570 or 650 (1:3000) and 4 drops of HRP blocker. Washing in between these solutions was performed twice at RT for 10 min with 0.2X SSCT prewarmed at 40 °C. Samples were incubated in primary antibody rabbit a-GFP (Invitrogen Cat. No. A11122) diluted (1:750) in blocking solution (0.1% PBTween, Normal goat serum 5% and 1% DMSO,1:750:) overnight at 4 °C then washed 3x with 0.1% PBTween for 1 hour with agitation. Samples were incubated in secondary antibody Goat a-Rabbit Alexa Fluor488 (Invitrogen Cat. No. A32731TR) diluted (1:750) in blocking solution for 3 hours at RT and then washed 6x 30 min with 0.1% PBTween. Samples were counterstained with 2 drops of DAPI provided in kit for 3 minutes and then rinsed once with 0.1% PBTween. Samples were mounted in 50% glycerol/PBS in glass bottom petri dishes. *phox2b* expression was quantified by manually counting the number of dots in each cell.

### Statistical analysis

All statistical analysis were carried out using GraphPad Prism 9. Normality of every sample was tested using the d’Agostino & Pearson test, followed by the Shapiro-Wilk test. Samples with a normal distribution were either compared using unpaired two-tailed t-test for samples with similar SD, Welch’s t test or Brown-Forsythe and Welch ANOVA tests for sample with different SD. Those without a normal distribution were compared through a Mann-Whitney U test. For all analyses, P values of under 0.05 were deemed statistically significant, with ****P<0.0001, ***P<0.001, **P<0.01, and *P<0.05.

## Results

### TNC migratory identity predicts their final fate

To define the relationship between migratory identity and differentiation outcome *in vivo*, we tracked single TNC from early stages until these reached their final destinations. This analysis shows that the movement of the chain is coherent, with cells maintaining relative positions as they move. In consequence, as leaders reach their destination and halt migration all the cells in the chain stop advancing and retain their positions (Figure 1A-B; Movie S1), comparable to a conga line where each member stops and remains in place as the first person stops. Leader cells initiate migration in an antero-posterior wave. TNC arising from segment 7-10 start movement at 15-16hpf and reach their target location 10-12h later. Leader cells divide perpendicularly to the direction of movement giving rise to a proximal and a distal daughter (Figure 1A-B (and E; Movie S1). The leader’s distal daughter remains at the front of the chain and reaches the most ventral location, lateral to the dorsal aorta, where it undergoes several rounds of divisions and forms the SCG. Neuronal differentiation is clear at the SCG location by 3dpf when strong expression of the noradrenergic marker *phox2bb* is detected (Figure 2A-C). Using the highly sensitive RNAscope technique we found that *phox2bb* is already strongly enriched in leader cells at early stages of migration (Figure 2I-L; Movie S2). Upon leader’s division, its proximal daughter the LP cell, arrests movement immediately dorsal to the SCG location (Figure 1A-B, Movie S1) where it envelopes the motor axon tracks and acquires a characteristic tubular morphology, differentiating into SC (Figure 2D). Follower 1 (F1), which migrates directly behind the leader, is the only cell in the chain that gives rise to more than one derivative, integrating the DRG and/or giving rise to SCs (Figure 1C-D, F and Figure 2D-H; Movie S3). Followers 2 and 3 (F2 and F3), which migrate at a more proximal position within the chain, advance only a short distance before coalescing ventral to the neural tube and exclusively populate the DRG (Figure 1C-D and F; Movie S3). Differentiation of DRG cells is first apparent by morphological changes at 30hpf, when cells show smaller sizes and a low cytoplasmic/nuclear ratio. By 36hpf the most lateral DRG cell initiates neural differentiation, increasing neurogenin 1 and reducing Sox10 expression. These changes are clearly observed by 3 and 5dpf (Figure 3A-F). Other DRG cells differentiate as glia as shown by expression of *foxd3* (8dpf; Figure 3G-I).

**Figure 1:**
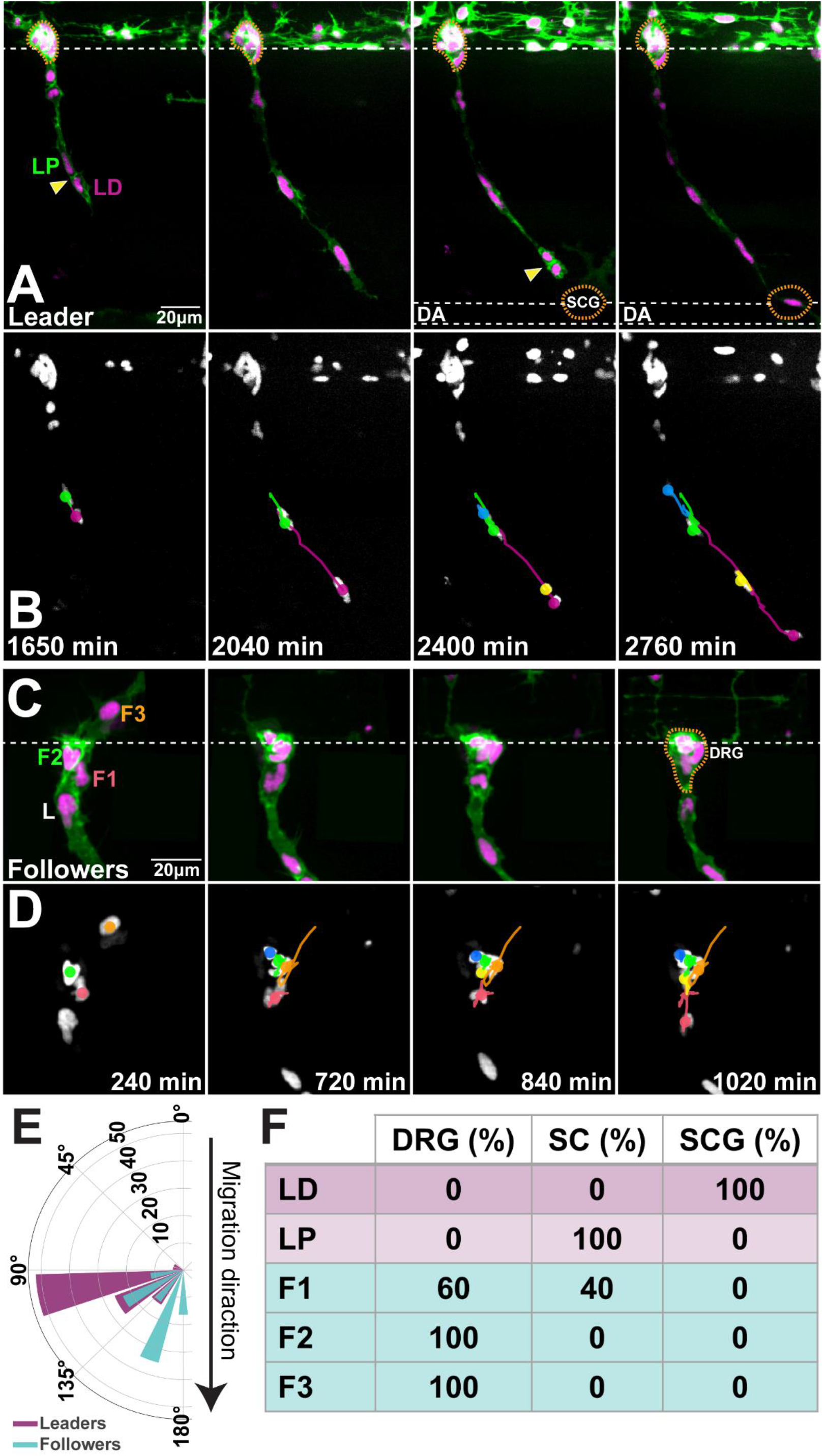
Migratory position predicts differentiation outcome. **(A, B)** Selected frames of **(A)** live imaging and **(B)** cell tracks between 2-3dpf of a Sox10:mG embryo at the level of somite 20. **(C, D)** Selected frames of **(C)** live imaging and **(D)** cell tracks of Sox10:mG between 22-36hpf at the level of somite 9. L: leader, F1: first follower, F2: second follower, F3: third follower. Orientation of the division plane (leaders n=27, followers n=43). **(F)** Derivatives formed by each cell in the migratory chain (n= 14 chains). Anterior to the left, dorsal top. Yellow arrows indicate divisions; orange dotted line DRG and SCG positions. LD leader’s distal and LP leader’s proximal daughters. DRG: dorsal root ganglia; SCG: sympathetic chain ganglia; SC: Schwann cell. Time in minutes after 18hpf.

**Figure 2:**
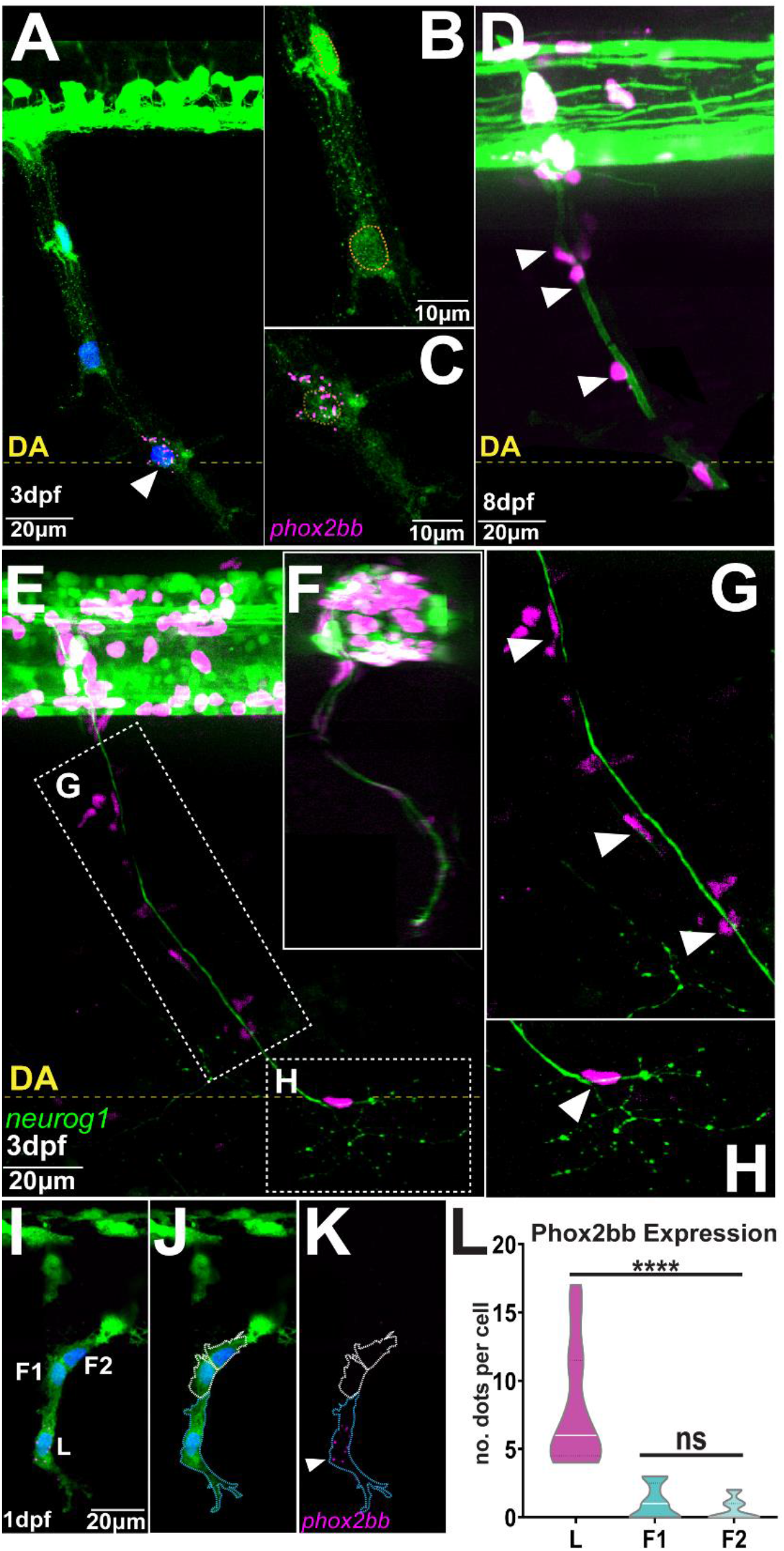
Phox2bb is predominantly expressed in leader cells. **(A-C)** *phox2bb* expression at 3dpf **(B)** Enlargements of follower and **(C)** leader cells. Sox10 in green, *phox2bb* in magenta and nuclei in blue in panel **A** and highlighted with orange dotted lines in **C-D. (D)** Confocal image of Sox10:mG embryo at 8dpf. White arrowheads indicate TNC cells showing Schwann cell morphology. **(E-H)** Confocal images of Sox10:mG; neurogenin1:GFP at 3dpf showing sensory neuron axon extension from DRG to SCG lateral view in **(E)** and transversal section in **(F). (G)** Enlargement showing TNC followers localising around the axon in position of Schwann cells indicated by white arrowheads and **(H)** Enlargement showing single leader cell in the position of SCG. **(I-K)** Confocal image of Sox10:GFP embryo at 1dpf showing expression of *phox2bb* in leader cell. L: Leader, F1 and F2: first and second followers. White arrowheads indicate leader. Blue and white dotted lines mark the outlines of leader and follower cells respectively. **(L)** Quantification of *phox2bb* expression (n=14 chains; Welch’s t test, L vs F1 and F2 p<0.0001, F1 vs F2 p=0.2566).Anterior to the left, dorsal top. DA: Dorsal Aorta.

**Figure 3:**
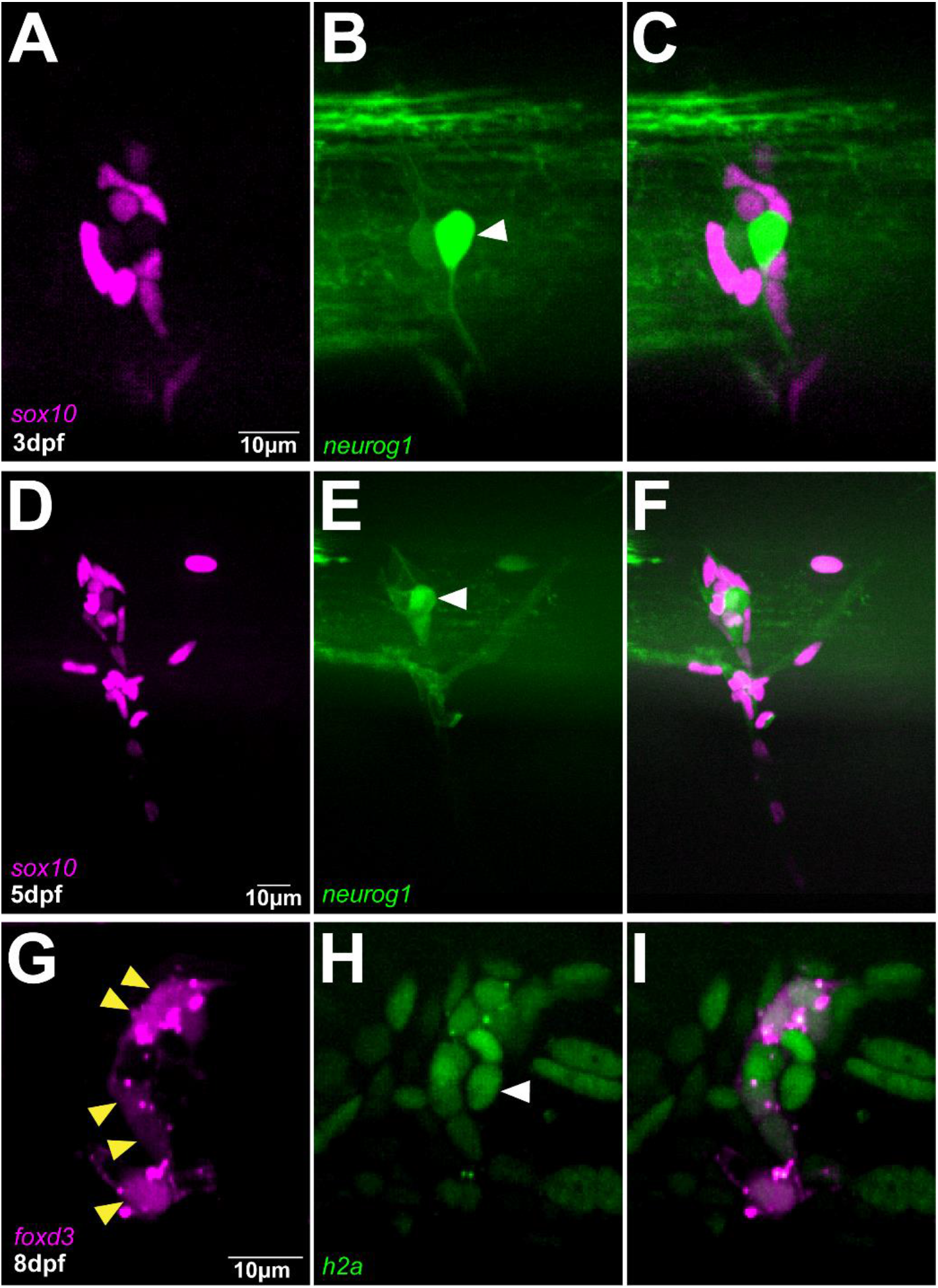
DRG is exclusively formed by follower. **(A-C)** Confocal images of Sox10:mG; neurogenin1:GFP at 3dpf showing the differentiation of a follower cell into the primary neuron of DRG indicated by white arrowhead. **(D-F)** Confocal images of in Sox10:mG; neurogenin1:GFP embryo at 5dpf showing a more mature DRG. **(G-I)** Confocal images of FoxD3:mCherry; H2aFVA:H2A-GFP embryo at 8dpf showing the peripheral followers expressing FoxD3, yellow arrowheads, and the primary neuron marked by the absence of FoxD3 expression, white arrowhead.

Taken together these results show that early migrating TNC are a coherent group in which the position of the cells in the migratory chain correlates with their fates. Interestingly, leader cells express the noradrenergic transcription factor *phox2bb* from early stages of development, suggesting that TNC migration is coordinated with fate specification. Moreover, our data show that the leader’s division results in cells of two fates, with only the distal cell giving rise to the SCG and the proximal daughter differentiating as a SC.

### Leaders TNC undergo asymmetric cell division that separates migratory identity

Our data show that upon division the leaders’ daughters adopt different fates. Hence, we hypothesised this mitotic event to be an asymmetric cell division. First, we analysed whether the leaders’ daughters present differences in size. *In vivo* imaging shows that the mitotic plane is positioned at the rear of the cell (Figure 4A) leading to an unequal partition of the cytoplasm upon cytokinesis (Figure 4B). Measurements of the cell area and volume show that the LD cell is 30% bigger than its proximal sibling (average area LD:130 ± 28 μm^2^, LP: 99 ± 23 μm^2^; Figure 4A and E; Movie S4; average volume LD: 1473 ± 125 μm^3^, LP: 876 ± 90 μm^3^; Figure 4B and F-G). This is in stark contrast to the followers’ divisions that is parallel to the direction of migration (Figure 1E) and where the cytokinesis plane bisects the cell into siblings of similar sizes (average area of 93 ± 26 μm^2^ and 90 ± 25 μm^2^; average volume 980 ± 160 μm^3^ and 987 ± 154 μm^3^; Figure 4C-F; Movie S4). Not only the leaders’ daughters show different sizes, but these cells acquire different migratory identities. The LD cell at the front of the chain retains the leader’s identity, moving faster (Figure 4H) and more directionally (Figure 4I) than the LP cell, which migrates as a follower (Figure 4H-I). The allocation of distinct migratory identities to the leader’s daughters may arise from the asymmetric distribution of determinants upon division, or it may be established through cell-cell interaction between the siblings only after division. In the former case differences in migratory behaviours will be evident immediately after division, while on the latter differences will only be observed sometime after division, with a lag period required for cells to interact and establish new identities. Tracking analysis shows that the LD cell moves faster and more directionally just 20 min after division, presenting the characteristic behaviour of a leader cell (LD cell directionality ratio 0.34 ± 0.10, speed 17.7 ± 4.3 μm/h; LP cell directionality ratio 0.27 ± 0.11, speed 14.2 ± 4.3 μm/h; Figure 4H-I), strongly suggesting that upon division the leaders’ daughters differentially inherit determinants that define their migratory identity. Next, we measured the directional correlation, which compares the actual direction of a cell path to the ideal direction of migration (a straight vertical vector). Positive values indicate cells migrate ventrally in a directed manner, values close to zero indicate the lack of directionality, while negative values signify cells that displace in the opposite direction, retracting dorsally. Leaders’ Distal daughters exhibited higher positive directionality (0.1045 ± 0.09; Figure 4J), while proximal daughters show poor directionality with a slightly negative value, indicating this cell retracts dorsally after division (−0.0091 ± 0.10; Figure 4J). Taken together, our results show that follower cells divide symmetrically giving rise to daughters with similar characteristics. On the other hand, leaders’ division is an asymmetric event, where the leaders’ distal daughter retains the leaders’ traits, while the proximal cell shows characteristics of a follower. Interestingly, the difference in migratory behaviour is observed with minimal temporal delay suggesting that migratory identity is segregated upon division.

**Figure 4:**
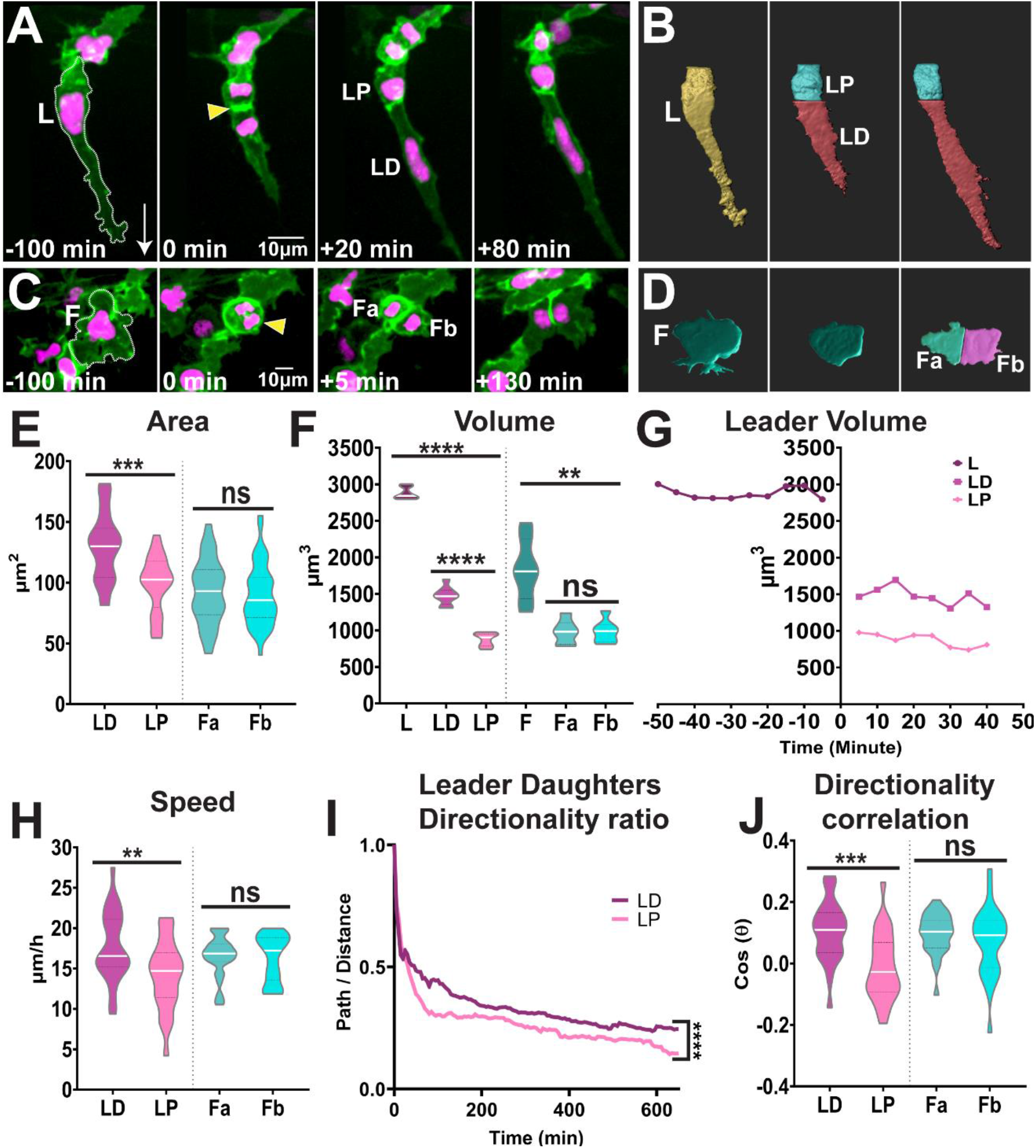
Leaders TNC divide asymmetrically while followers divide symmetrically. **(A-B)** Confocal images of Sox10:mG **(A)** and 3D rendering **(B)** of leader asymmetric division into distal and proximal daughters. L: Leader, LD: Leader Distal, LP: Leader Proximal; white arrow indicates direction of migration and yellow arrowhead points to the dividing cell. **(C-D)** Confocal images of Sox10:mG **(C)** and 3D rendering **(B)** of follower symmetric division. F: Follower, Fa and Fb Follower’s daughters. **(E)** Quantification of cell area of leaders’ and followers’ daughters quantified immediately after division (n=25 for leaders and n=44 for followers; Unpaired t test, p=0.0001 for LD vs LP and p=0.5852 for Fa vs Fb). **(F)** Quantification of cell volume of leaders, followers and their daughters after division (n= 10 divisions for leaders and n=10 divisions for followers; Brown-Forsythe and Welch ANOVA tests, p<0.0001 for L vs LD and LP, p<0.0001 for LD vs LP, p=0.0012 for F vs Fa and Fb, p>0.9999 for Fa vs Fb). **(G)** Cell volume over time of a representative leader before and after division. **(H)** Quantification of speed of movement of leaders’ and followers’ daughters (n= 25 for leaders and n=44 for followers; Unpaired t test, p=0.0072 for LD vs LP, p= 0.8690 for Fa vs Fb). **(I)** Directionality ratio of leaders’ daughters (n= 25 divisions; Simple linear regression, p<0.0001 for LD vs LP). **(J)** Directionality correlation of leaders’ and followers’ daughters (n= 25 for leaders and n=44 for followers; Welch’s t test, p=0.0004 for LD vs LP, p=0.2798 for Fa vs Fb).

## Discussion

TNC invading the medial pathway migrate collectively forming single file chains that depend upon cells adopting distinct identities. Leader cells, at the front of the chain, direct the movement of the group. Followers, form the body of the chain and track leaders. Herein we show that the cells in the chain maintain their relative positions during migration and these correlate with their final destination and terminal differentiation. Consistent with previous reports (Dutton et al., 2001; Raible et al., 1992; Raible and Eisen, 1994), our data show that the majority of follower cells give rise to single derivatives. *In vivo* imaging shows that leader cells populate the ventral-most locations, lateral to the dorsal aorta, and give rise to the SCG clonally, confirming earlier observation from fixed specimens (An Min et al., 2002). Followers divide symmetrically forming two cells of similar sizes that maintain their migratory identity, which may contribute to their similar differentiation outcome. Leader cells on the other hand, divide asymmetrically and perpendicular to the direction of movement, giving rise to a larger distal daughter that retains the leader’s identity and a smaller proximal sibling that migrates as a follower. The difference in migratory behaviour between siblings is observed almost immediately after division, strongly suggesting that migratory identity is allocated by the unequal distribution of determinants. In other developmental contexts, such as bristle patterning in *Drosophila*, the asymmetric positioning of the mitotic spindle ensures the unequal inheritance of Notch components, which in turn defines the cell identity and subsequent differentiation programme (Lu and Johnston, 2013; Schweisguth, 2015). Interestingly, Notch components such as Numb have been shown to be asymmetrically inherited during division of avian TNC (Wakamatsu et al., 2000). Moreover, Notch activity controls the expression of signalling components that orchestrate the mitotic spindle polarity and orientation, α-adaptin and Numb (Charnley et al., 2020). Hence, it is possible that the asymmetric distribution of Notch signalling controls identity allocation during the leader’s division and maintains a single leader cell per chain, akin to its role in identity allocation before migration initiation (Alhashem et al., 2021a). The asymmetric division of the leader cells is also reminiscent of tip cell divisions during blood vessel formation (Costa et al., 2016). Tip cells position their spindle towards the rear of the cell resulting in siblings of different sizes and morphology that present distinct migratory identities. In this case the unequal partition of VEGF signalling components (*kdlr* mRNA and ERK activity) is responsible for migratory identity allocation. Whether a comparable mechanism is at play in TNC requires further study.

Our data show that the expression of the transcription factor *phox2bb* is enriched in leader cells defining it as a leader marker and posing interesting questions about the specification state of TNC during migration. *phox2bb*, orthologue of the human *PHOX2B*, encodes a paired like homeobox required for sympathetic and enteric neuron development. It is expressed by NC as these reach their target sites (Ernsberger et al., 1995; Pei et al., 2013) and its absence (knock-out or morphant embryos) prevents the differentiation of the enteric and sympathoadrenal linages (Elworthy et al., 2005; Ernsberger and Rohrer, 2018; Pattyn et al., 1999; Pei et al., 2013; Taylor et al., 2016). Our use of a new and more sensitive gene expression techniques has revealed *phox2bb* expression much earlier during TNC development, prompting the proposition of the new model of NC fate acquisition (Kelsh et al., 2021). The cyclical fate restriction model explains the seemingly paradoxical fact that NC maintain multipotency and at the same time express differentiation markers of one or more fates. It proposes that NC are highly dynamic progenitor cells that move between unstable sub-states which are temporarily biased towards one of the possible differentiation outcomes. This transient differentiation bias is likely defined by the expression of key fate determination receptors, while the final fate decision depends on the time spent in a particular state and the level of activity of the determination receptor. The model also proposes that the ‘dwell-time’ in any of these biased sub-states can be altered by local environmental inputs. As migration concludes, follower cells forming the body of the chain remain exposed to neuronal axons, in an environment enriched with differentiation cues for SC, while followers closer to the neural tube, such as F3, are exposed to sensory neuronal and glial specification cues, explaining the relationship between migratory identity, migration chain position and the differentiation outcome observed in our study.

Our observations of persistent *phox2bb* expression in leader cells is consistent with them being at least strongly biased towards sympathetic neuronal fate. The asymmetric division of leader cells may unequally distribute fate determination receptors and/or activity, such as Notch (see above), hence defining a distinct differentiation bias for each of the sibling cells. Furthermore *phox2bb* has been shown to regulate Notch receptor expression (Revet et al., 2008), and our data show that Notch signalling in turn controls *phox2bb* transcription in TNC (Alhashem et al., 2021a). Hence, it is possible that Notch and *phox2bb* establish a positive forward loop that reinforces each other’s expression only in the leader cell. As the leader cell reach the dorsal aorta the expression of *phox2bb* is further reinforced by BMP signalling from this site, leading to the expression of sympathetic fate effectors (Müller and Rohrer, 2002; Rohrer, 2011; Saito et al., 2012; Schneider et al., 1999). Further experimental analysis and mathematical modelling is required to define the role of such a possible Notch-Phox2bb-BMP circuit during TNC differentiation.

## ACKNOWLEDGMENTS

We are especially grateful to N. Daudet for his scientific and personal support. To the KCL fish facility staff. This project was funded by MRC G1000080/1, Royal Society 2010/R1 and Wellcome Trust 207630/Z/17/Z to CL; and BBSRC BB/S015906/1 to RNK.

## AUTHOR CONTRIBUTIONS STATEMENT

ZA and CL designed the research. ZA and KCS performed the experiments. CL and RNK directed the work. CL wrote the manuscript.

## Supplementary movies

**Movie S1: Leaders’ distal daughter resides at the position of SCG**.

**(A)** Fluorescent images and **(B)** cell tracks of Sox10:mG Time-lapse imaged between 2dpf and 3dpf showing the leader’s daughters migrating ventrally and dividing, with the Distal daughter residing and dividing at the position of the SCG suggesting a clonal origin of the SCG. Yellow arrowheads indicate the points of division, L: Leader, LD: Leader Distal, LP: Leader Proximal, nuclei in magenta or grey and membranes in green, time in minutes past 18hpf. Related to Figure 1.

**Movie S2: *phox2bb* is exclusively expressed in leaders**.

3D projection of Sox10:GFP at 24hpf showing expression of *phox2bb* leader TNC. Neural crest membranes in green, nuclei in blue and *phox2bb* RNA in magenta. Related to Figure 2.

**Movie S3: Followers give rise to the DRG**.

**(A)** Fluorescent images and **(B)** cell tracks of Sox10:mG time-lapse imaged between 22hpf and 36hpf showing followers coalescing lateral to the neural tube to from the DRG. Yellow arrowheads indicate the points of division, nuclei in magenta or grey and membranes in green, L: leader, F1: first follower, F2: second follower, F3: third follower, time in minutes past 18hpf. Related to Figure1.

**Movie S4: Leaders divide asymmetrically, while followers divide symmetrically**.

**(A)** Time-lapse of Sox10:mG imaged between 22hpf and 25hpf showing a leader cell maintaining its elongated shape and dividing asymmetrically into two daughters with different sizes. **(B)** Time-lapse of Sox10:mG Imaged from 20hpf to 23hpf showing a follower cell rounding up and dividing symmetrically into two daughters with similar sizes. yellow arrowheads indicate divisions, L: Leader, LD: Leader Distal, LP: Leader Proximal, F: Follower, Fa and Fb: Follower daughters, neural crest membranes in green, nuclei in magenta, time in minutes. Related to Figure 4.

